# Inference of tissue relative proportions of the breast epithelial cell types luminal progenitor, basal, and luminal mature

**DOI:** 10.1101/2021.05.17.444493

**Authors:** Thomas E. Bartlett, Peiwen Jia, Swati Chandna, Sandipan Roy

## Abstract

Single-Cell Analysis has revolutionised genomic science in recent years. However, due to cost and other practical considerations, single-cell analyses are impossible for studies based on medium or large patient cohorts. For example, a single-cell analysis usually costs thousands of euros for one tissue sample from one volunteer, meaning that typical studies using single-cell analyses are based on very few individuals. While single-cell genomic data can be used to examine the phenotype of individual cells, cell-type deconvolution methods are required to track the quantities of these cells in bulk-tissue genomic data. Hormone receptor negative breast cancers are highly aggressive, and are thought to originate from a subtype of epithelial cells called the luminal progenitor. In this paper, we show how to quantify the number of luminal progenitor cells as well as other epithelial subtypes in breast tissue samples using DNA and RNA based measurements. We find elevated levels of cells which resemble these hormone receptor negative luminal progenitor cells in breast tumour biopsies of hormone receptor negative cancers, as well as in healthy breast tissue samples from *BRCA1* (*FANCS*) mutation carriers. We also find that breast tumours from carriers of heterozygous mutations in non-BRCA Fanconi Anaemia pathway genes are much more likely to be hormone receptor negative. These findings have implications for understanding hormone receptor negative breast cancers, and for breast cancer screening in carriers of heterozygous mutations of Fanconi Anaemia pathway genes.

## 1 Introduction

Despite advances in diagnosis and treatment in the last several decades, breast cancer remains responsible for more than 5000 deaths per year in the U.K. [2]. Triple-negative breast cancer (TNBC) is a particularly aggressive subtype of breast cancer, with poor prognosis and few treatment options [3]. The triple-negative breast cancer subtype has this name because it lacks receptors for the hormones estrogen and progesterone, as well as the growth-factor HER2. Hence, quantifying hormone receptor negative cells in breast tissue that is at risk of developing cancer is likely to provide important prognostic information. The cell of origin for TNBC is thought to be a hormone receptor negative epithelial cell called the ‘luminal progenitor’ cell [4]. Luminal progenitor cells are one of three main cell subtypes of the breast epithelium, along with hormone receptor positive luminal mature cells, and basal epithelial cells: these cells are arranged in luminal and basal layers in the bi-layered epithelium of the breast. Luminal progenitor cells, basal cells, and luminal mature cells are all thought to originate from mammary stem cells, which also reside in the basal layer [5].

Cellular identity is set during human development [6, 7], and is corrupted in disease [8]. In this article, we show that cells-of-origin of aggressive breast cancers [4] may be tracked in bulk tissue genomic data, a data-type which is scaleable to much larger cohorts than single-cell technologies. Single-Cell Analysis has revolutionised genomic science in recent years. However, due to cost and other practical considerations, single-cell analyses are impossible for studies based on medium or large patient cohorts. For example, a single-cell analysis usually costs thousands of pounds for one tissue sample from one volunteer, meaning that typical studies using single-cell analyses are based on very few individuals. While single-cell genomic data can be used to examine the phenotype of individual cells, cell-type deconvolution methods are required to track the quantities of these cells in bulk-tissue genomic data.

DNA methylation (DNAme, Fig.1) is an epigenetic mark that is applied to the DNA at specific cytosine bases, and DNA methylation patterns enable different tissues and cell types to be distinguished from one another [9, 1]. DNA methylation in bulk-tissue samples is typically analysed in terms of the DNA methylation rate *β* at each DNA cytosine base. Hence, the methylation rate *β* represents the fraction of DNA strands in the bulk sample (from several hundred thousand cells) which carry the methylation mark at a specific cytosine locus. Because DNAme marks are chemical modifications of specific DNA cytosine bases, their average levels over a tissue or cell type fluctuate much more slowly than levels of transient mRNA transcripts do. The relative stability, and the tissue and cell type specificity of DNAme marks, along with the observation that variation in mRNA levels may account for less than half of the variation in protein concentrations [10, 11], means that DNAme data are a promising alternative to gene-expression or RNA-seq data for identifying cell type mixing proportions in bulk-tissue data. DNAme patterns also have a lot of potential as prognostic biomarkers for various cancers, as changes in DNAme levels in a tissue precede formation of cancer at the same site [12]. DNA methylation patterns have been shown to have prognostic power in TNBC [13] as well as other womens’ cancers [14, 15].

**Figure 1:**
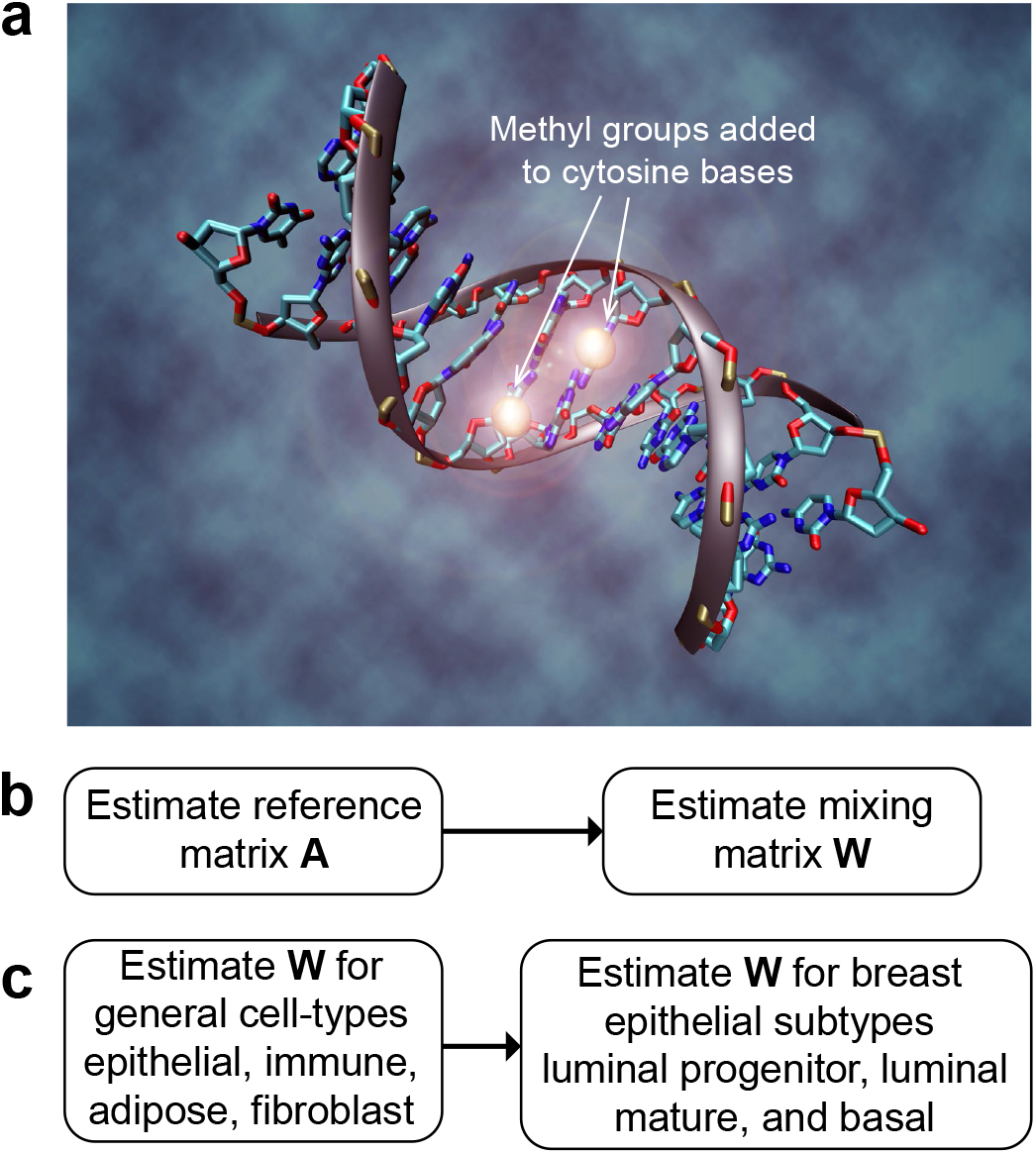
(a) DNA methylation: a methyl group is added to some cytosine DNA bases, giving tissue and cell-type specific patterns at the genome-wide scale that allow accurate inference of cell-type proportions in bulk-tissue genomic (i.e., convolved) data. (b) We use a two-step procedure to first estimate the reference matrix **X**, before second estimating the mixing matrix **W** (representing the cell-type proportions). (c) DNAme data are used to estimate general cell-type proportions with an earlier tool [1], before DNAme and RNA-seq data are used to estimate breast epithelial subtype proportions. Illustration by Christoph Bock, Max Planck Institute for Informatics.

The basic form of the cell-type deconvolution model is

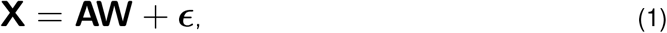

where **X** ∈ ℝ^*p*×*n*^ are the observations on *p* genomic features in *n* bulk-tissue (i.e., mixed cell-type) samples, **A** ∈ ℝ^*p*×*k*^ are the genomic profiles over the *p* features in *k* cell types, and **W** ∈ [0, 1]^*k*×*n*^ are the mixing weights representing the *k* cell or tissue-type proportions in the *n* samples. Various methods exist to estimate matrix **W** for gene-expression (microarray or RNA-seq) data **X** after estimating matrix **A** [16, 17, 18, 19]. The reference matrix **A** can be thought of as representing a *p*′-dimensional space in which the cell-types of interest can be distinguished well from one-another. In practice, **A** may be determined as part of a unified procedure for fitting the model of Eq.1 [20, 21], or **A** may be estimated *a-priori* from external reference data-sets [1, 22, 16, 18, 19]. In this paper we focus on inference of **W** using *a-priori* estimates of **A**. Estimating **A** *a-priori* typically consists of estimating the expected value ℝA_*il*_ based on data from a specific feature (e.g., gene) *i* in a specific cell-type *j*, for features *i* ∈ {1, …, *p*′} and cell-types *l* ∈ {1, …, *k*}. These *p*′ features are typically identified by differential expression analysis for each cell type individually (in comparison with all the other cell types pooled) [23]. In this article we propose an alternative based on the Mahalanobis distance (i.e., to take account of redundancy due to correlated features/genes).

In this work, we show how to estimate the concentrations of the breast epithelial cell subtypes luminal progenitor, luminal mature and basal. We do this using both DNA methylation and RNA-seq (i.e., gene-expression) data. We note that for RNA-seq data it is challenging to accurately estimate the relative proportions of breast adipose cells using the available reference data for general adipose cells. Hence for RNA-seq data we estimate the epithelial subtypes as a proportion of the epithelial compartment only. Whereas for DNAme data, we estimate the epithelial subtypes as a proportion of all relevant cell-types (also including adipose and stromal). We show the robustness of our DNAme-based classifier, and evaluate it against several RNA-based methods in terms of estimation of epithelial subtype proportion. We then show that in the presence of a wide range of cell-types that our DNAme-based breast epithelial subtype classifier reproduces expected proportions of the epithelial subtypes in biopsies of hormone receptor positive and negative tumours. Using our DNAme-based breast epithelial subtype classifier, we explain important differences in inter-individual tumour heterogeneity in terms of variations in luminal progenitor cells that are associated with carriers of heterozygous mutations in non-BRCA Fanconi Anaemia pathway genes.

## 2 Results

In this section, we first present the results of a simulation study; we then compare inferences of cell type proportions of breast epithelial cell subtypes based on data from DNA and RNA; finally, we present our findings when applying these methods to data from breast tumour biopsies and healthy breast tissue samples. We note that ideally, we would evaluate methodology with respect to a known ground-truth data-set. However, we also note that accurate solutions to the deconvolution problem are tissue-specific: e.g., in the human breast it is challenging to identify epithelial subtypes in the presence of fat. There is currently no data-set with known ground-truth for evaluating deconvolution methodology designed for the breast epithelial subtypes. Instead, we compare epithelial cell-type proportions inferred in bulk-tissue genomic data from different experimental platforms on the same samples. By comparing inferences of cell-type proportions derived from DNA (i.e., DNAme) with those derived from RNA (i.e., mRNA), we provide confidence in the inferences available from our proposed methodology with a surrogate validation that approximates validation with a ground-truth data-set.

### Simulation study

For our DNAme reference profiles for the breast epithelial subtypes luminal progenitor, luminal mature, and basal, we used previously-published DNAme data from bisulphite-sequencing experiments [24], leading to reference matrix **A** ∈ [0, 1]^58×3^ (where *A*_*il*_ ∈ [0, 1] rather than *A*_*il*_ ∈ ℝ because we also have methylation rate *β* ∈ [0, 1], for further details see Methods). To estimate **W** (the relative proportions of the breast epithelial subtypes), we used a hierarchical procedure, first estimating the relative proportions of the general cell types epithelial, adipose, fibroblast, and immune [1], before estimating the proportions of the breast epithelial subtypes. We estimate the tissue relative proportions *W*_:,*j*_ of *k* cell-types in bulk-tissue sample *j* as:

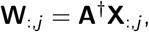

where † is a pseudo-inverse of **A** calculated with a robust linear model using the using the `rlm` () function of the `MASS` package in `R` [1]. Hence, we refer to **A**^†^ calculated in this way as the RLM-pseudo-inverse (RLM-PI) of reference matrix **A**. The elements of **W**_:,*j*_ must be constrained to sum to 1, and furthermore in practice, this robust linear model fit may lead to some small negative values that are assumed to be noise. We apply an adaptive noise threshold, such that any values as great in magnitude as the most negative value are replaced with 0.

To test how robust the procedure is for estimating concentrations of luminal progenitor, luminal mature and basal epithelial subtypes from DNAme data, we generated simulated data based on Eq.1, together with this pre-defined breast epithelial subtype reference matrix **A**, with *n* = 100 simulated bulk-tissue samples, as follows. After generating 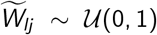 independently for *l* ∈ {1, 3}, and *j* ∈ {1, 100}, and using this to calculate the simulated 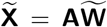, we then replaced some randomly-chosen instances of these simulated data values 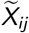 (where *i* and *j* are independently sampled) with random noise 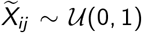. We repeated this procedure *b* = 1000 times. Figure 2 shows the results of these robustness tests. We find that with up to 40% of data-values replaced with noise, the results remain very good.

**Figure 2:**
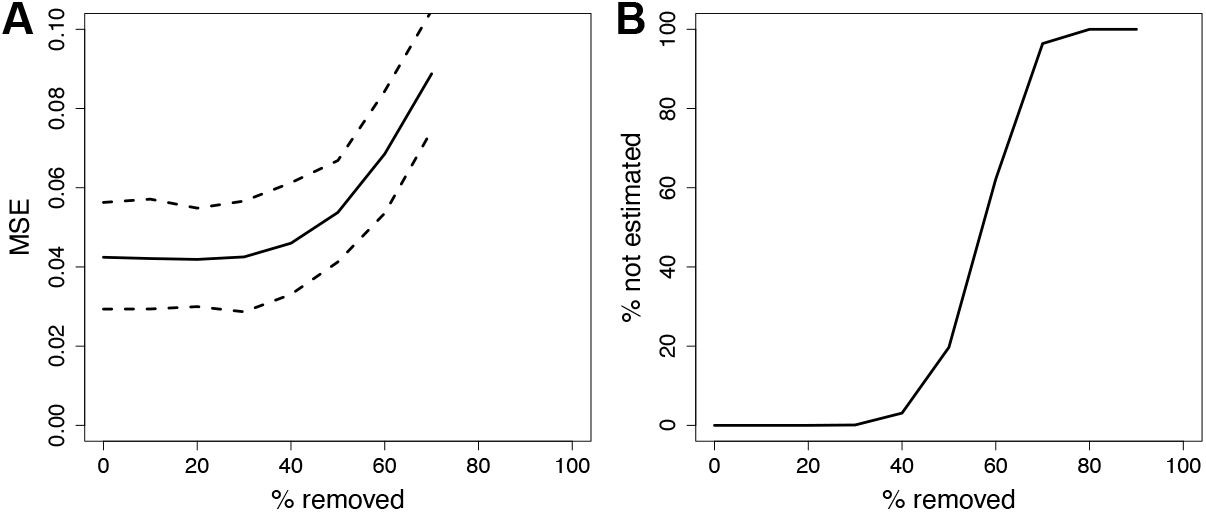
Robustness of inference. A data-set with n = 100 bulk-tissue samples was simulated, and data-points were removed and replaced at random with noise; this was repeated b = 1000 times. (a) Mean square error (MSE) comparing predicted proportions of epithelial subtypes with the ground-truth, as an increasing number of data-points are removed. Dashed lines indicate 95% C.I.s. (b) Percentage of mixing proportions l ∈ {1, k}, W_lj_, j ∈ {1, n}, which cannot be estimated, as an increasing number of data-points are removed.

### Comparison of RNA and DNA based inference of epithelial subtypes

For RNA-seq reference profiles for the breast epithelial subtypes luminal progenitor, luminal mature, and basal, we used publicly-available single-cell RNA-seq data for *n* = 13909 breast epithelial cells [5]. We identified which cells corresponded to each breast epithelial subtype, by fitting a Gaussian mixture model (GMM) in the Eigenspace of the graph-Laplacian [25], a procedure we refer to as GMM-LE clustering. To implement GMM-LE clustering, we carry out degree-corrected regularised spectral clustering [26] replacing the *k*-means clustering step by fitting a Gaussian mixture model. Fig.3 shows UMAP (uniform manifold approximation and projection) [27] representations of the data-matrix in two dimensions, illustrating clusters detected by GMM-LE, and average expression levels of independent marker genes for the breast epithelial subtypes luminal progenitor, luminal mature, and basal.

**Figure 3:**
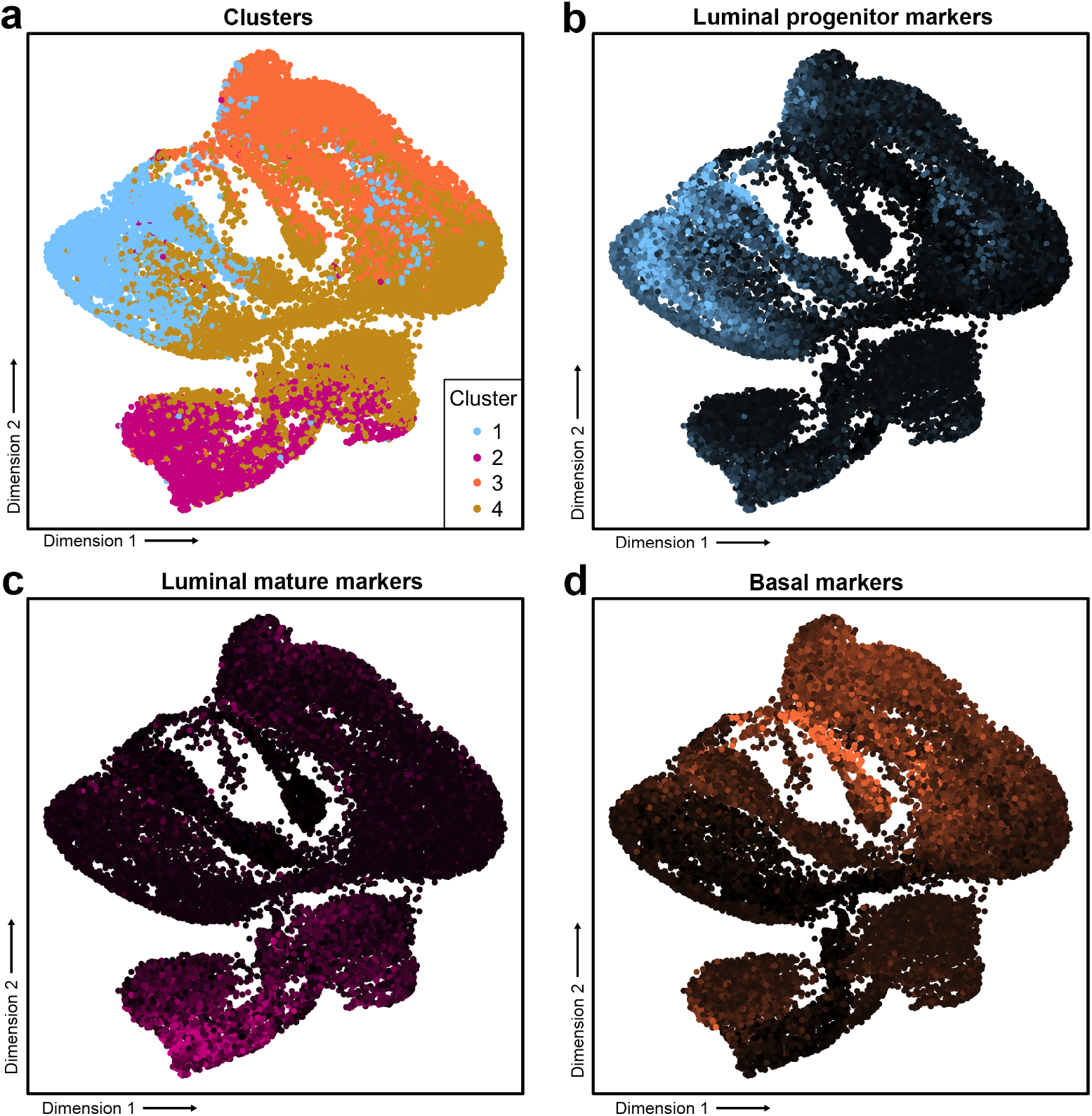
UMAP projections of single-cell RNA-seq data from n = 13909 breast epithelial cells. (a) Inferred GMM-LE clusters. Colour intensity matching shows strong overlap of (a) with (b)-(d) in which colour intensities correspond to mean log expression levels from independent and pre-defined sets of marker genes for luminal progenitor, luminal mature, and basal cells (respectively).

We compared the tissue proportions of the breast epithelial subtypes luminal progenitor, luminal mature and basal, inferred from DNAme with those inferred from RNA-seq data according to a range of existing methods, based on matched data from *n* = 175 low-stage (I-II) breast tumour biopsies downloaded from TCGA (The Cancer Genome Atlas). We did this as a surrogate for evaluation against a known ground-truth data-set, instead evaluating breast epithelial subtype proportions inferred in bulk-tissue genomic data from different experimental platforms on the same samples, i.e., by comparing inferences of cell-type proportions derived from DNA with those derived from RNA. Table 1 shows correlation coefficients, comparing inferred proportions of the breast epithelial cell subtypes estimated from DNAme data, with those estimated from RNA-seq data according to RLM-PI as well as the URSM [16], Bisque [18], MuSiC [17] and Cibersortx [19] methods. We also used two methods for determining which genes/features to include in our *a-priori* estimate of matrix **A** (full details are given in Section 4). Briefly, method (1) uses a statistic based on the Mahalanobis distance (i.e., it reduces redundancy in the chosen space of **A** by adjusting for covariance) to rank features / genes (Table 1a). Alternatively, method (2) uses a test-statistic for each feature/gene obtained by testing for differential expression using the `edgeR` package in `R` (Table 1b). In both cases, the genes/features are ranked based on the extent to which they discriminate each cell-type from all the others pooled (with each cell-type taking turns). Noting again that the correlation of mRNA levels with levels of their corresponding proteins is often around 0.4 [10, 11], i.e., when comparing expression levels quantified by different genomic data modalities, the results shown in Table 1 show good agreement when estimating **W** with the RLM-PI method, after using method (1) to estimate **A** (Section 4). Furthermore, the wide variety of correlation coefficients shown in the results of Table 1 indicate the importance of choosing an appropriate method for estimating matrix **A** (i.e., compare Table 1a with Table 1b), as well as in estimating matrix **W** (i.e., compare within Table 1a and Table 1b). We note that all comparisons are between epithelial cell-types. This is due to the difficulty for RNA-seq data to accurately estimate the relative proportions of breast adipose cells using the available reference data for general adipose cells. Hence, we only estimate relative proportions of epithelial cell subtypes using the RNA-seq data, whereas we estimate these epithelial cell subtypes as well as the relative proportions of adipose and stromal cells when using DNAme data.

**Table 1:** Correlation coefficients (with 95% C.I.s) compare estimated epithelial cell-type proportions from bulk-tissue genomic data obtained simultaneously from the same samples via DNAme and mRNA measurements according to various methods designed for mRNA data, in n = 175 low-stage breast tumour samples. (a) and (b) show results using matrix **A** estimated using two approaches which are labelled ‘method 1’ and ‘method 2’ respectively, as described in Section 4 below.

| **a** | Method | Luminal progenitor | Luminal mature | Basal |
| --- | --- | --- | --- | --- |
|  | RLM-PI | 0.88 (0.84-0.91) | 0.86 (0.81-0.89) | 0.52 (0.41-0.62) |
|  | URSM [16] | -0.65 (-0.72-0.55) | 0.08 (-0.06-0.23) | 0.46 (0.34-0.57) |
|  | Bisque [18] | 0.53 (0.41-0.63) | 0.51 (0.39-0.61) | 0.12 (-0.03-0.26) |
|  | MuSiC [17] | 0.49 (0.37-0.59) | 0.52 (0.41-0.62) | 0.22 (0.07-0.35) |
|  | Cibersortx [19] | 0.85 (0.81-0.89) | 0.86 (0.82-0.89) | 0.51 (0.39-0.61) |
| **b** | Method | Luminal progenitor | Luminal mature | Basal |
|  | RLM-PI | 0.08 (-0.07-0.22) | 0.13 (-0.02-0.27) | 0.23 (0.09-0.37) |
|  | URSM [16] | 0.46 (0.33-0.57) | 0.71 (0.62-0.77) | 0.39 (0.26-0.51) |
|  | Bisque [18] | 0.45 (0.32-0.56) | 0.59 (0.48-0.68) | 0.25 (0.1-0.38) |
|  | MuSiC [17] | 0.57 (0.47-0.67) | 0.56 (0.45-0.65) | 0.24 (0.1-0.38) |
|  | Cibersortx [19] | 0.41 (0.27-0.52) | 0.24 (0.09-0.37) | 0.14 (-0.01-0.28) |

### Application to breast cancer biopsies and controls

Luminal progenitor cells are thought to be the cell-of-origin for triple-negative breast cancer (TNBC), which fits well with the model of luminal progenitor cells being similarly hormone receptor negative (HR-) [4]. To test whether the proportions of the tumour cells with similar genomic characteristics to the breast epithelial subtypes luminal progenitor, luminal mature, and basal (as inferred from DNAme data) are in line with this biological model, we applied our DNAme breast epithelial cell type deconvolution method to DNAme data from TCGA (The Cancer Genome Atlas). Fig.4a shows the concentrations of the breast epithelial subtypes estimated from DNAme data in hormone receptor positive and negative cancers (as classified by expression levels of ESR1, PGR and ERBB2/HER2 in matched gene expression microarray data). As well as elevated levels of luminal progenitor-like cells in HR-(hormone receptor negative) breast cancer, Fig.4a also shows elevated level of luminal mature-like cells (luminal mature cells are HR+) in HR+ breast tumours, also in line with this biological model. To further test whether the proportions of the breast epithelial subtypes inferred from DNAme data are in line with established biological theory, we analysed a publicly-available data-set from GEO (Gene Expression Omnibus). It is well known that the breast epithelial tissue of heterozygous carriers of *BRCA1* mutations contains higher levels of luminal progenitors than would be expected in the general population [28], and this is confirmed in Fig.4b. *BRCA1* (*FANCS*) mutation carriers who unfortunately go on develop breast cancer mostly develop tumours which are HR-[28], presumably due to inefficient DNA damage repair, which is also in line with this model.

**Figure 4:**
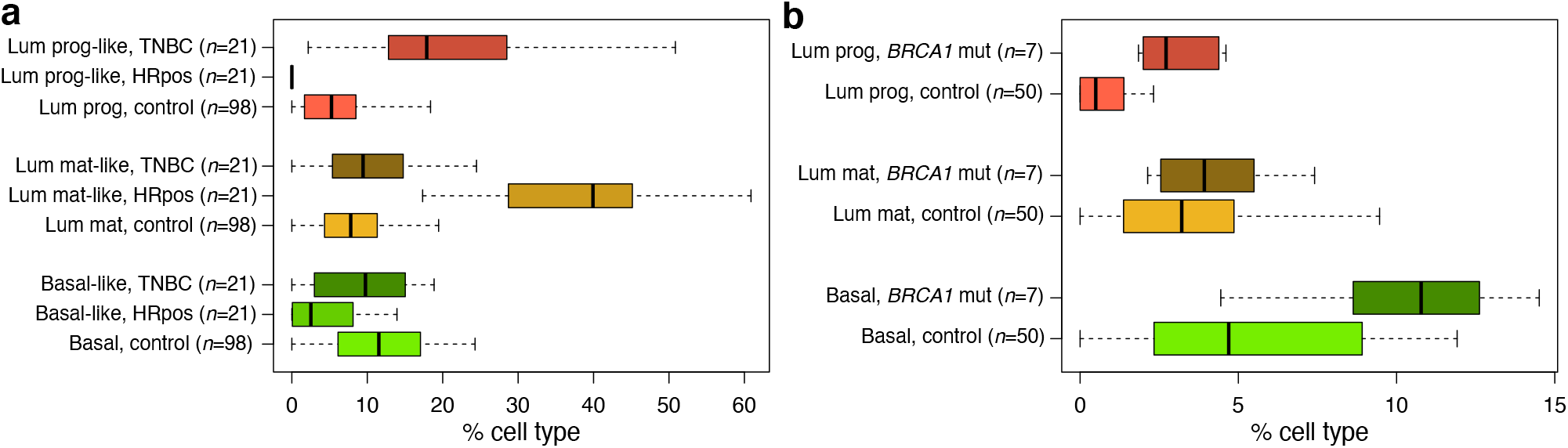
Inferred proportions of luminal progenitor (lum prog), basal, and mature luminal (lum mat) cells, in (a) hormone receptor positive / negative cancers. We use ‘like’ as a suffix to indicate similar genomic characteristics of tumour cells and their related healthy epithelial cells. (b) Inferred proportions of luminal progenitor, basal, and mature luminal cells, and in (b) BRCA1 (FANCS) mutation carriers. Inferences are based on DNAme data.

The *BRCA1* (*FANCS*) mutation is the best known example of a mutation in the Fanconi Anaemia (FA) / DNA damage repair pathway. Next, we investigated whether similar observations could be made in non-BRCA FA-pathway genes (non-BRCA FANC genes). We used DNAme data from *n* = 257 tumour samples from TCGA with matched clinical data to test how heterozygous carriers of mutations in non-BRCA FANC genes affect the proportions of cells with genomic characteristics of the breast epithelial subtypes luminal progenitor, luminal mature, and basal cells in tumours that unfortunately arise in this population. To do this, we used multivariate linear models to explain the variation in the concentrations of luminal progenitor, luminal mature, and basal-like cells in tumour biopsies in terms of non-BRCA FANC gene mutation status, as well as the clinical covariates age and disease stage. We observed that carriers of non-BRCA FANC genes have significantly elevated levels of luminal progenitor-like and basal-like cells (Fig.5a-b, significance indicated by C.I. bars), but significantly decreased levels of mature luminal-like cells, as well as of epithelial cells overall (Fig.5c-d, epithelial proportions calculated as the sum of the proportions of the individual epithelial subtypes). This again fits with the biological model of inefficient DNA damage repair in heterozygous carriers of mutations in FA pathway genes, with this manifesting as elevated levels of cells with replicative potential in the epithelial tissue (i.e., luminal progenitors and basal cells). Presumably this pool of cells with replicative potential is prone to expand more quickly in this population who may have inefficient DNA damage repair mechanisms due to heterozygous mutations in FA pathway genes. This may be compounded by somatic mutations that arise by chance in the other allele of the gene with the FA pathway mutation. This would be expected to correspond to an increase in prevalence of HR-tumours relative to HR+ tumours in FA-gene mutation carriers, as previously reported [29], and this is confirmed in Fig.5e-f. The 10 HR-tumour biopsies from carriers of heterozygous mutations in non-BRCA FA genes represented in Fig.5e correspond to 2 carriers with SNPs in each of FANCD2, FANCF, FANCG, FANCI, and FANCM (one carrier had SNPs in both FANCG and FANCI), as well as one carrier with a SNP in FANCB. Several of these genes have already been associated with breast cancer susceptibility; particularly, FANCD2 [30], FANCG [31, 32], and FANCM [32].

**Figure 5:**
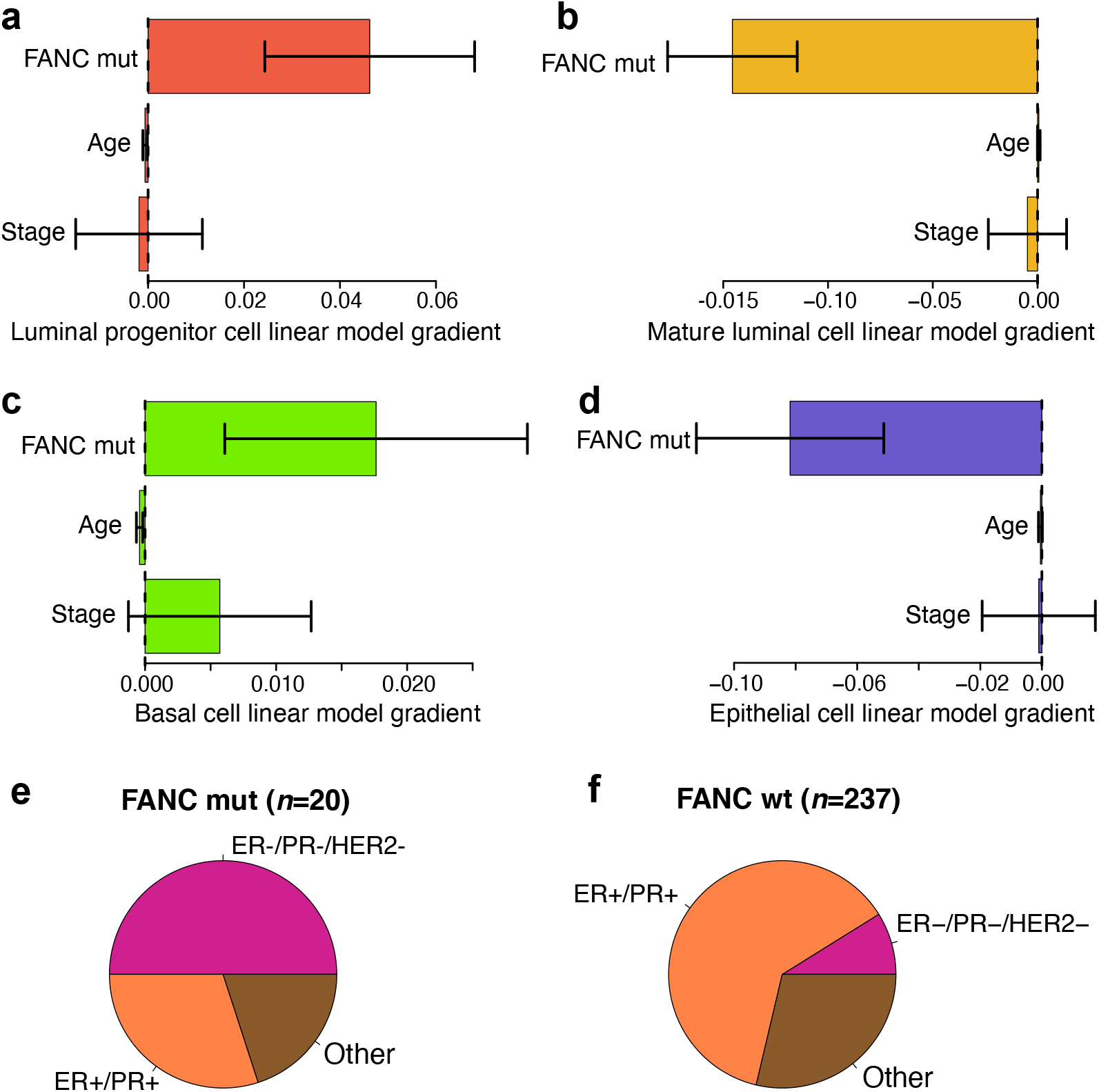
Linear modelling (with 95% C.I.s indicated) of estimated proportions of (a) luminal progenitor-like, (b) mature luminal-like, (c) basal-like, and (d) overall epithelial-like cells, in n = 257 tumour biopsies. The horizontal bars arranged along the vertical axis in each plot represent linear model predictors as follows: FA-gene (non-BRCA, FANC suffix) mutation carrier status, disease stage, and volunteer age. (e) Biopsies of FA-gene mutation carriers are mostly hormone receptor (HR) triple-negative, whereas those of (f) FA-gene wild-type are mostly HR-positive. Inferences are based on DNAme data.

## 3 Discussion

We have presented methods to infer proportions of the breast epithelial cell subtypes luminal progenitor, luminal mature, and basal, using DNA and RNA-based methods. We have compared the results using several existing algorithms for estimating the matrix of mixing proportions **W** ∈ [0, 1]^*k*×*n*^, for *k* cell types in *n* bulk-tissue samples. In particular, we found good agreement (as assessed by Pearson correlation) between our DNAme (DNA methylation) based inference method and our RNA-seq based inference method when estimating the mixing matrix **W** using RLM-PI (pseudo-inverse calculated via robust linear modelling), after estimating the profile matrix **A** using our Mahalanobis distance based statistic for ranking features/genes (Section 4). The best correlations between inferences from different data modalities are much stronger than the previously-reported level of agreement between expression levels of a gene quantified by different genomic data modalities. However we note that this study could be improved with access to an accurate ground-truth data-set. Future work will also include refining the proposed methodology to consider more epithelial subtypes determined by more subtle distinctions, and identified by further development of our proposed GMM-LE unbiased clustering / cell-type discovery methodology.

We note that for RNA-seq data it is challenging to accurately estimate the relative proportions of breast adipose cells using the available reference data for general adipose cells. Hence for RNA-seq data we estimate the epithelial subtypes as a proportion of the epithelial compartment only. Whereas for DNAme data, we estimate the epithelial subtypes as a proportion of all relevant cell-types (also including adipose and stromal). Our results when applying these methods to tumour biopsies and healthy breast tissue samples agree well with established biological models of the cell-of-origin for hormone receptor negative breast cancers [4]. Specifically, we found significantly elevated levels of luminal progenitor-like cells (luminal progenitors are hormone receptor negative) in biopsies of triple-negative (i.e., hormone receptor negative) breast cancers. Furthermore, we found significantly elevated levels of luminal progenitors in the healthy breast epithelial tissue of heterozygous carriers of *BRCA1* (*FANCS*) mutations, as previously reported [28].

Investigating whether these findings extend to non-BRCA FA-pathway genes (non-BRCA FANC genes), we found that the concentration of luminal progenitors in a tissue sample was statistically significantly predicted by the presence of a heterozygous non-BRCA FANC mutation, in a model that adjusts for the important clinical covariates disease stage and patient age. This is in line with observations based on the same data-set, that incidence of hormone receptor negative tumours make up a significantly much greater proportion of the total number of tumours in heterozygous carriers of non-BRCA FANC mutations (50% compared to 9%). This finding has implications for cancer screening in this population of heterozygous carriers of non-BRCA FANC mutations [33], given the relative much worse prognosis of hormone receptor negative breast tumours compared to breast cancers overall, and may also be relevant to screening for other epithelial cancers in this population.

## 4 Methods

### Estimating matrix A

We wish to estimate **A** ∈ ℝ^*p*′×*k*^ for *p*′ < *p* ‘characteristic features’ which define a space which best separates the *k* cell-types of interest, optimised as follows. Define **Y** ∈ ℝ^*p*×*m*^ to be the observations on the same *p* genomic features in *m* single-cell samples over the *k* cell-types of interest (as in Fig.1).Then, for each *k* select *p*′ < *p* dimensions such that

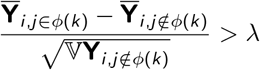

for all *i* ∈ {1, …, *p*′} (noting a reordering with respect to *i* ∈ {1, …, *p*}), where *ϕ* (*k*) is the set of cells of type *k*, and 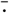 and 𝕍 are the mean and variance operators. This can then be refined by choosing the *p*′ dimensions that maximise a modified Mahalanobis distance:

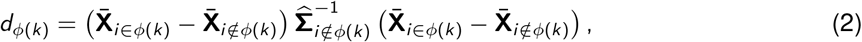

where 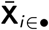 is the vector of mean values in each dimension *j* ∈ {1, …, *p*} over the single-cells of certain type(s), and 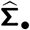 is the corresponding data-estimate of the covariance matrix. Noting that Eq.2 can be factorised into two identical factors

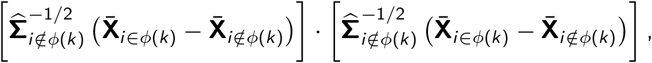

the distance in Eq.2 can be optimised by choosing the *p*, dimensions (for each *k*) that have the largest values in the vector

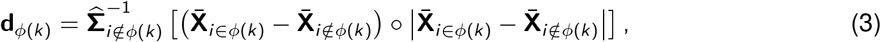

where ○ represents element-wise multiplication. The matrix **A** was estimated from average expression levels of independent marker genes for the breast epithelial subtypes luminal progenitor, luminal mature, and basal. These class labels were identified using the GMM-LE clustering, then used to define **A** ∈ ℝ^*p′*×*k*^, with the *p*′ dimensions chosen either by method (1) according to Eq.3, or by method (2) which consists of identifying the most significantly upregulated genes for each cell type *l* ∈ {1, …, *k*}, when compared to all other cell types by using the `edgeR` package in `R` (with default settings) to test genes for differential expression. The dimensionally-reduced space was then defined by the union of the lists of top 250 ranked genes / transcripts for each cell-type, according to either method (1) or method (2).

### DNAme data processing

Bulk-tissue DNAme microarray data were downloaded from TCGA (https://www.cancer.gov/about-nci/organization/ccg/research/structural-genomics/tcga) and GEO (https://www.ncbi.nlm.nih.gov/geo/) and pre-processed as follows. Data were background-corrected, probes were removed with < 95% coverage across samples, and any remaining probes with detection *p* > 0.05 were replaced by *k*-NN imputation, with *k* = 5. Cell type specific bisulphite-sequenced DNAme data for the breast epithelial subtypes luminal progenitor, luminal mature, and basal were downloaded from EGA (European Genome-phenome Archive), https://ega-archive.org) and pre-processed as follows. Sequencer reads were aligned and counted using Bismark [34] with default settings. We subsequently retained only reads mapping to DNA cytosine loci that are also measured in the DNAme microarray data, and which had a total number of mapped reads (methylated+unmethylated) of at least 20 (leading to a granularity of methylation rate of at least 0.05). In this breast epithelial cell type specific data-set, only one reference profile is available for each epithelial subtype. So to select *p*′ features (i.e., DNA cytosine loci) of interest, we selected features according to the following criteria: (1) Low variance of methylation rate *β* across cell types of non-epithelial lineages (specifically, 𝕍*β <* 0.001, where 𝕍 is the variance operator). (2) A difference in mean methylation rate *β* of at least 0.5 between the epithelial subtype of interest and non-epithelial lineages. (3) The greatest difference in mean methylation rate *β* between the breast epithelial subtypes. This results in a reference matrix **A** ∈ [0, 1]^58×3^. To estimate the concentrations of the breast epithelial subtypes in the presence of stromal and adipose cells, we use a hierarchical procedure, first estimating the relative proportions of the general cell types epithelial, adipose, fibroblast, and immune [1], before estimating the proportions of the breast epithelial subtypes. Linear modelling of breast epithelial subtype proportions was carried out with subtype proportion as the response and FA-gene mutation carrier status, disease stage, patient age as predictors using the `lm()` function in `R`.

### RNA-seq data processing

Single-cell RNA-seq data for breast epithelial cells were downloaded from GEO (https://www.ncbi.nlm.nih.gov/geo/), and libraries with at least 250 reads and transcripts expressed in at least 1000 cells were analysed further. Batch effects were removed using the COMBAT software [35]. These data were used to define our RNA-seq reference profiles for the breast epithelial subtypes. To estimate the concentrations of these epithelial subtypes in the presence of stromal and adipose cells, we augmented this single-cell breast epithelial data-set with synthetic data based on bulk RNA-seq libraries from purified breast stromal [24] cells and from adipose tissue samples from ENCODE (Encyclopedia of DNA Elements, https://www.encodeproject.org). We carried out this augmentation by randomly sampling a single cell library from the breast epithlelial data-set, then replacing the values of the chosen adipose or stromal bulk library with those of the equivalent quantiles of the sampled breast epithelial single cell library, as follows. Define 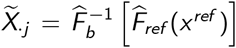, where *X*^*ref*^ ∈ ℝ^*p*^ is the vector of (adipose or stromal) reference values across all *p* genes, 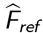 is the empirical CDF of *X*^*ref*^,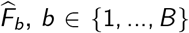 is the equivalent empirical CDF of a randomly selected epithelial cell, and 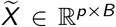 is the data-matrix of sampled, normalised values for the adipose or stromal reference to use in the subsequent cell type deconvolution procedure. Default settings were used for all deconvolution methods, including URSM (`Python` implementation), Bisque and MuSiC (`R` packages), and Cibersortx (online web tool). GMM-LE clustering took place in the `R` software environment using the `svds` () function from the `rARPACK` package to do the Eigendecomposition of the graph Laplacian (to obtain the ‘Laplacian Eigenspace’, LE), and the `Mclust ()` function from the mclust package to fit the Gaussian mixture model (GMM), with cells then assigned to clusters according to the maximum posterior probabilities. Default settings for all packages were used unless otherwise specified.

### Availability of data and materials

The breast epithelial cell subtype bulk-data deconvolution tool is made available as an R package from https://github.com/tombartlett/BreastEpithelialSubtypes

DNA methylation microarray data for breast cancer and controls were downloaded from TCGA and from GEO under accession number GSE69914. Matched gene-expression microarray data for breast cancer and controls were downloaded from TCGA.

Bisulphite-sequenced DNAme data for the purified breast epithelial cell subtypes luminal progenitor, luminal mature, and basal, were downloaded from the European Genome-Phenome Archive (EGA) under accession number EGAS00001000552

Single-cell RNA-seq data for the breast epithelial cell subtypes luminal progenitor, luminal mature, and basal, were downloaded from the Gene Expression Omnibus (GEO) under accession number GSE113197

Bulk RNA-seq data for breast stromal cells were downloaded from the European Genome-Phenome Archive (EGA) under accession number EGAS00001000552. Bulk RNA-seq data from the adipose tissue samples ENCFF072HRK, ENCFF654JLY, ENCFF732LRY and ENCFF924FNY were downloaded from ENCODE.

## 5 Additional information

### Author contributions

TB conceived of and designed the study and wrote the manuscript. All analyses were carried out by TB and PJ with advice from SC and SR.

### Funding

The work of TB during this project was funded by the MRC grant MR/P014070/1. The funding body had no role in the design of the study, collection, analysis, and interpretation of data, or writing of the manuscript.

## Acknowledgements

TB dedicates his work on this manuscript to the memories of Joanne Walker and Joel Walker. The authors are grateful to Peter Kennedy for reading the manuscript and for his feedback which has improved the work. The results published here are in whole or part based upon data generated by The Canadian Epigenetics, Epigenomics, Environment and Health Research Consortium (CEEHRC) initiative funded by the Canadian Institutes of Health Research (CIHR), Genome BC, and Genome Quebec. Information about CEEHRC and the participating investigators and institutions can be found at http://www.cihr-irsc.gc.ca/e/43734.html

## Competing interests

The authors declare that there are no competing interests.

## Notes

### Competing Interest Statement

The authors have declared no competing interest.

### Summary of Updates

Revisions to the paper have been implemented, following feedback from colleagues.

